# Optoregulated force application to cellular receptors using molecular motors

**DOI:** 10.1101/2020.03.31.015198

**Authors:** Yijun Zheng, Mitchell K.L. Han, Renping Zhao, Johanna Blass, Jingnan Zhang, Dennis W. Zhou, Jean-Rémy Colard-Itté, Damien Dattler, Markus Hoth, Andrés J. García, Bin Qu, Roland Bennewitz, Nicolas Giuseppone, Aránzazu del Campo

**Author notes:** This authors contributed equally to this work (Yijun Zheng, Mitchell K.L. Han, Renping Zhao, Johanna Blass).

## Abstract

Mechanotransduction events in physiological environments are difficult to investigate, in part due to the lack of experimental tools to apply forces to mechanosensitive receptors remotely. Inspired by cellular mechanisms for force application (i.e. motor proteins pulling on cytoskeletal fibers), here we present a unique molecular machine that can apply forces at cell-matrix and cell-cell junctions using light as an energy source. The key actuator is a light-driven rotatory molecular motor linked to polymer chains, which is intercalated between a membrane receptor and an engineered biointerface. The light-driven actuation of the molecular motor is converted in mechanical twisting of the polymer chains, which will in turn effectively “pulls” on engaged cell membrane receptors (integrins, cadherins…) within the illuminated area. Applied forces have the adequate magnitude and occur at time scales within the relevant ranges for mechanotransduction at cell-friendly exposure conditions, as demonstrated in force-dependent focal adhesion maturation and T cell activation experiments. Our results reveal the potential of nanomotors for the manipulation of living cells at the molecular scale and demonstrate, for the first time, a functionality which at the moment cannot be achieved by any other means.

---

External mechanical stimuli are sensed and translated by cells into biochemical signals in a process called mechanotransduction^1,2^. In turn, biochemical signals regulate cellular and extracellular mechanical properties. This mechano-sensitive feedback modulates cellular functions as diverse as proliferation, differentiation, migration and apoptosis, and is crucial for tissue formation, homeostasis, repair and pathogenesis. Perturbations along the chain of mechanical sensing and biochemical responses lead to various pathological disorders such as loss of hearing, cardiovascular dysfunction, muscular dystrophy and cancer. Being able to apply mechanical signals to cells at the molecular scale in cell culture models or tissues would be of great value in the understanding of these diseases.

Cells generate forces at the molecular scale through directional polymerization of actin filaments and by the action of myosin as ATP-driven molecular motor^3^. The objective of the present study rests on the application of external forces to cells by using an artificial, purely synthetic molecular motor^4^. Among synthetic molecular devices, light-driven rotary motors based on over-crowded alkene molecules are able to autonomously cycle unidirectional 360° rotations around their central double bond by using light and temperature as combined sources of energy^5^. When individual rotary motors are coupled to fixed elements through pairs of polymer chains, the rotational actuation of the motor enforces conformational twisting of the polymer chains. In this configuration, the original rotary motion is transformed into a linear motion, with the potential to pull on individual nanoscale objects^4,6–9^. By making use of this actuation principle, we envisioned using such a motor-polymer conjugate to apply external mechanical forces to individual membrane receptors of a living cell, and to trigger mechanotransduction processes therefrom.

The general molecular design used for the present study includes a light-driven molecular motor with a so-called stator part (in blue) and a rotor part (in red) (Figure 1a). The stator part is linked to two triethylene glycol (TEG) chains connected to the membrane receptor of interest (e.g. integrin or T cell receptor) via a complementary ligand (i.e. RGD or antiCD3). The rotor part is linked to two flexible polyethylene glycol (PEG) chains (M_w_ ≈ 5000 g.mol^-1^) and is connected to a synthetic interface (hydrogel) by covalent bonds. The orthogonal chemical protocols used for the two coupling steps *(i.e.* catalyzed and non-catalyzed click reactions) are selective and, therefore, no cross-reactions are expected (details in SI). In this configuration, light irradiation progressively rotates the motor that twists the two pairs of polymer chains and effectively shortens the receptor-interface link. As a consequence, a tensional force is applied directly on the membrane receptor, pressumably engaged with the contractile cytoskeletal machinery. Previous studies have demonstrated that the energy produced by a single motor in a similar polymer conjugate is in the range of 12 kT (or 50 pN·nm), with a typical frequency of rotation in the range of 1 to 0.01 Hz at room temperature^6,7^. This range of forces and time scales seems appropriate for applying forces to cells. This study demonstrates the implementation and application of this molecular tool in two relevant mechanotransduction scenarios: force-dependent focal adhesion maturation and force-dependent T cell activation.

**Figure 1.**
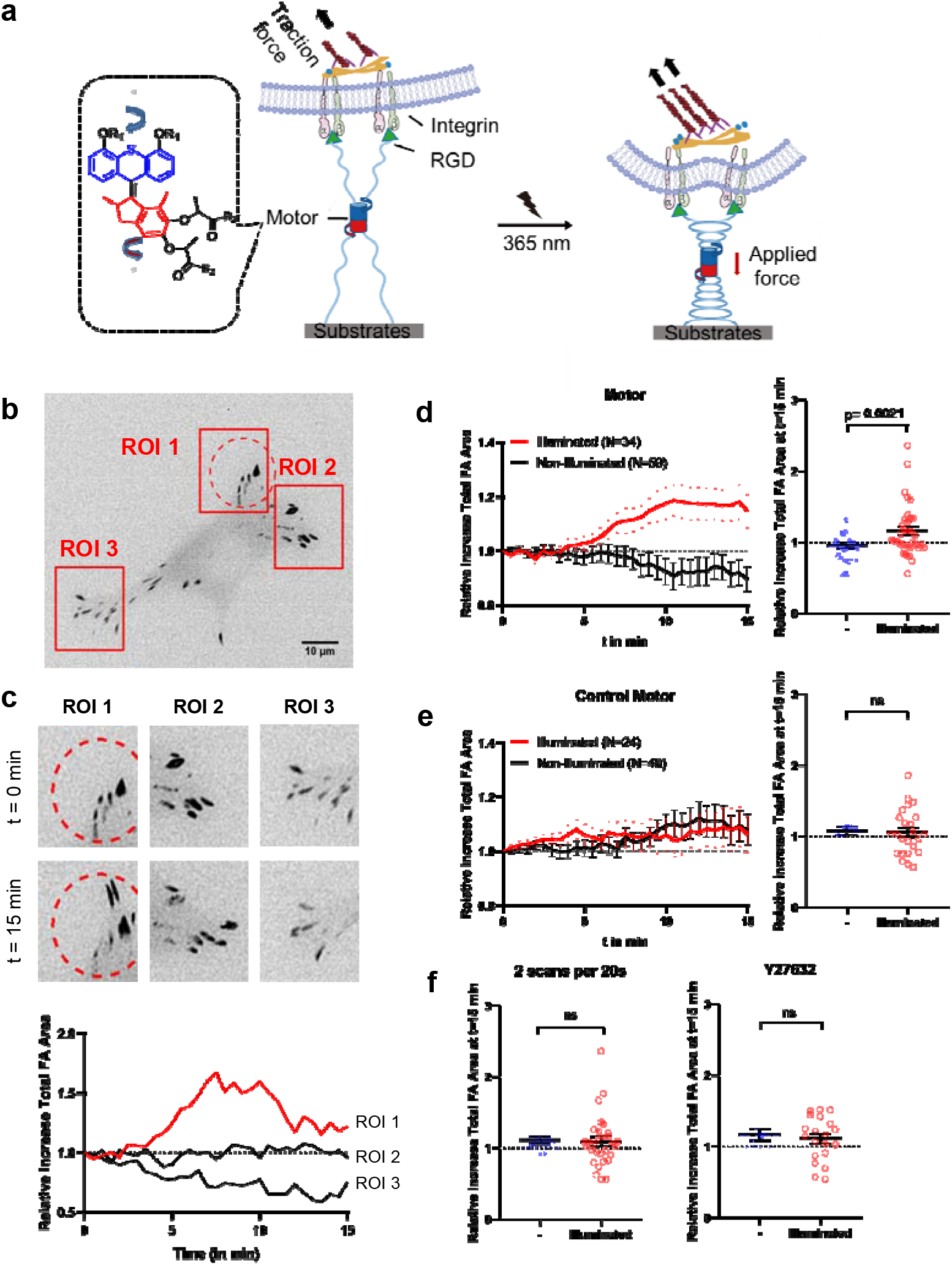
Light-driven force application by the motor/PEG/surface construct coupled to RGD integrin-binding ligands leads to focal adhesion growth and maturation. (a) Scheme showing the design of the force application platform with the rotary motor. (b) Representative image of focal adhesions of MEFs expressing paxillin-RFP adhering to the molecular motor-RGD substrate tracked over time during substrate activation using UV-light scans. The activated area is demarcated by the red dotted lines. (c) zoom-ins of the ROIs at time points t=0 min and 15 min (left) and the effect of substrate activation on the total area of Focal Adhesions was analyzed within defined ROIs (red boxes in b). (d) Plots represent the mean ± s.e.m. of relative increase in Total Focal Adhesion Area of analyzed within N# of ROIs of cells seeded on motor substrate illuminated with 5 scans per 20s. The scatterplot on the right represents the relative increase in Total FA Area at t=15 min with the result from each ROI plotted individually. ROIs were analyzed from 34 cells from six independent experiments. Nonilluminated ROIs (n=59), and illuminated ROIs (n=34). (e) Same as in (d) but cells were seeded on the control motor substrate. N# of ROIs from 24 cells from 5 independent experiments. Non-illuminated ROIs (n=40), and illuminated ROIs (n=24). (f) Scatterplots (mean ± s.e.m) with each datapoint representing an analyzed ROI at t=15 min. Cells were UV-illuminated with 2 scans per 20s (left – 30 cells from 3 independent experiments – Nonilluminated ROIs (n=59), and illuminated ROIs (n=30)) or with 5 scans per 20s but pretreated with ROCK-inhibitor Y27632 (right – 20 cells from three independent experiments – Non-illuminated ROIs (n=32), and illuminated ROIs (n=20)). Data from non-illuminated and illuminated areas within each condition were compared using a non-paired one-tailed Student’s t-test.

After optimization of the coupling reactions for specific and stable cell binding to the surface-immobilized ligand/motor/PEG conjugate (see SI for details and supplemental figures S1,S2), we first sought to regulate focal adhesion (FA) growth through force application at the RGD/integrin complex. Mouse Embryonic Fibroblasts (MEFs) expressing paxillin-RFP to mark FAs were seeded on the RGD/motor/PEG surface and allowed to adhere overnight. We illuminated circular areas (62 μm^2^) of the cells containing several FAs using a 365 nm point-by-point scanning laser. Exposure conditions were 5 scans every 20s for 15 min (see SI for details). We tracked the total FA area in the exposed and in control (not exposed) regions of the same cell by imaging the RFP-paxillin every 30 seconds for 15 minutes, a timescale which has been reported for FA growth after force application^10,11^. Fig. 1b shows an example of a MEF cell after illumination. The FA area in the illuminated region or the cell (ROI1, Fig. 1c) increased visibly with exposure, while the FA area in the control regions (ROI2 and ROI3, Fig. 1c) did not. Quantification of ROIs from multiple cells (for N see figure legends) showed, on average, a rise in total FA area in illuminated regions starting at 5 minutes, and an increase in Total FA Area of about 20% at 10 minutes, which is sustained until the end of the measurement. In contrast, the Total FA Area in non-illuminated regions from the same cells decreased to about 95% within the same timespan (Fig 1d), showing that illumination induced a significant difference in FA growth compared to FA growth in non-illuminated areas. To rule out that UV irradiation on its own could induce the observed FA area changes, similar experiments were performed in MEF cells on control surfaces conjugated with non-rotary motors (locked by a single episulfide moiety in place of the rotating double bond, Supplemental Scheme 1). In this case no difference in total FA area between UV-illuminated and non-illuminated regions was observed (Fig. 1e). A lower exposure dose (2 scans per 20s, i.e. 40% of the initial dose) did not result in measurable differences in FA area between illuminated and non-illuminated areas (Fig. 1f and Supplemental Fig 4a), indicating that the dose (and consequently force) was not enough to trigger a cellular response. Intermittent exposure programs (i.e. 3 min on – 3 min off – 5 runs per 20s) did not elicit any cellular response either (Supplemental Fig. 4b, c). Additionally, by perturbing myosin contractility via ROCK inhibition with Y27632 (10 μM), the observed motor-induced increase of the FA area was lost (Fig. 1f right and Supplemental Fig. 5), confirming the involvement of actomyosin contractility in the force-induced FA reinforcement and maturation^2^. Together, these results indicate that the RGD/motor/PEG surface was able to locally apply forces on integrin complexes within FAs upon UV-light exposure, leading to downstream integrin mechanotransduction and FA area increase^12^ in an illumination dose-dependent manner. Note that the observed responses and time scales are in agreement with experimental results from established methods in mechanotransduction research (see SI).

Mechanical stimulation on the T cell receptor (TCR) leads to T cell activation, as proven in single-molecule experiments^13–16^. We tested if the opto-mechanical actuator coupled to the T-cell receptor (TCR) was able to activate T cells by light exposure. For this purpose, Jurkat and primary T cells were loaded with Ca^2+^ indicator fluo-4-AM and seeded on antiCD3/motor/PEG modified surface (Fig 2a). CD3 is a key component of TCR for signal transduction, and thus Ca^2+^ influx was induced in the cells when they contacted the surface, indicating recognition of the anti-CD3 antibody. In order to decouple the Ca^2+^ influx induced by the motor from a possible response by simple interaction with the surface, Jurkat cells were seeded at 0 mM [Ca^2+^] concentration, and the same volume of 2 mM [Ca^2+^] was added at 8 mins post-seeding. At 15 min cells were exposed to pulses (1 s) of 365 nm light for 1 min. A Ca^2+^ rise was observed immediately after the first UV pulse (Fig. 2b, statistics in Fig. 2c), which decayed within 5 min. In comparison, no response was observed when the Jurkat cells were seeded on control motor (no rotation) surfaces (Figs. 2b and 2c). These results indicate that the observed response was not associated to UV illumination per se, but it was a consequence of the applied pulling force on the TCR by the rotary motor linked to antiCD3. Under the same conditions, primary human CD4^+^ T cells showed a similar response (Fig. 2d and 2e). Shorter light pulses (0.5 s) did not elicit an observable response (Fig. 2f and 2g), indicating that a certain threshold of force is required for force-mediated activation of T cells. Longer pulses (2 s) induced a decrease in the Ca^2+^ signal on the motor and control surfaces (Fig. 2h and 2i), which is most likely related to photodamage of the cells following long UV pulses. These results define the boundaries for experimentation with this unique tool. As negative control, similar experiments were performed with the CD28 receptor, whose activity has been proven to be unaffected by mechanical stimulation^17^. No response was observed (Supplemental Fig.7), confirming that the observed force-dependent Ca^2+^ response was specific to the CD3 engagement. Mechanical force applied by the motor does not change the contact area at the immunological synapse (IS) (Supplemental Fig.8). Together, these results demonstrate, for the first time, that synthetic molecular machines can be used to apply external forces to specific membrane receptors on T cells and study mechanotransduction events at biointerfaces. It should be noted that a similar motor had been recently incorporated to the membrane of living cells, and used to kill cells by drilling pores in the membrane upon 360 nm illumination.^18^

**Figure 2.**
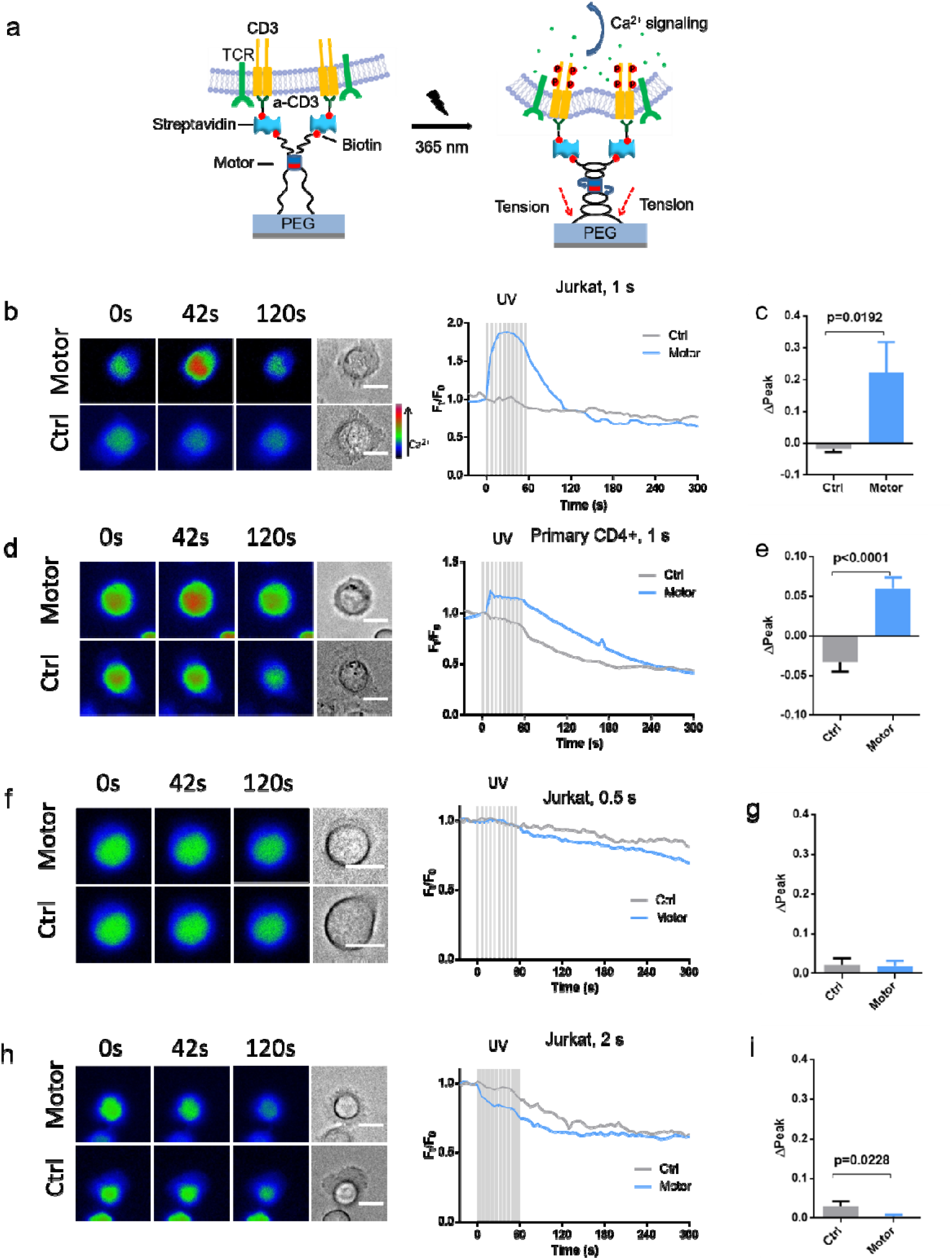
Mechanical force generated by rotatory motor can trigger TCR signaling. (a) Schematic of manufacture and activation of anti-CD3 antibody linked to the substrate via rotatable motor (Motor) or non-rotatable motor (Ctrl). Ten UV pulses were applied within 60 sec to activate the motor. (b-e) Ca^2+^ influx is induced by activation of the motor. Either Jurkat T cells (b,c,) or primary human CD4^+^ T cells (d,e) loaded with Fluo-4-AM were settled on motor substrate for 20 min prior to UV illumination (starting at time 0). Duration of UV pulses was 1 s. Exemplary cells and Ca^2+^ traces are shown in (b) and (d). Ca^2+^ influx (ΔPeak) was analyzed in c (Ctrl, n = 137 cells from 6 independent experiments; Motor, n = 153 cells from 9 independent experiments) and e (Ctrl, n = 42 cells from 2 independent experiments; Motor, n = 46 cells from 3 independent experiments). (f-i) Shorter or longer duration of UV pulses cannot induce Ca^2+^ influx. We used Jurkat cells and applied UV pulses with either shorter duration of 0.5 s (f,g, Ctrl, n = 119 cells from 5 independent experiments; Motor, n = 201 cells from 9 independent experiments) or extended duration of 2 s (h,i, Ctrl, n= 101 cells from 5 independent experiments; Motor, n = 172 cells from 9 independent experiments). Scale bars are 10 μm.

In order to quantify forces and pulling distances provided by our molecular motor construct, we tracked the light-induced pulling of microparticles tethered to motor-chain conjugates against the drag force of liquid flow in a microfluidic channel (Fig 3a). This method was inspired by experiments using centrifugal forces^19^. The optical quantification of the tethered microparticle motion requires spacers much longer than the PEG_5000_ linker used for the cell experiments. Here we used a dsDNA chain of 1.7 μm length. The 53-fold increase in chain length is counteracted by the 57-fold increase in persistence length, rendering the coiling and twisting of the DNA construct a scaled version of that of PEG5000 linker. Typical time-displacement curves for a microparticle are shown in Figure 3b, details of the experiments are given in SI. In the control experiment with the non-rotary motor, the bead was displaced by 1 μm in the flow direction during the first minute of flow, followed by continuous displacement (creep). UV exposure did not affect on the displacement of the beads connected by the non-rotary motor. In contrast, beads tethered to the surface via rotary motors were retracted against the drag force within the first minute of UV-irradiation. We quantified the light-induced length reduction as ratio of the negative displacement after 2 minutes of illumination to the expected bead displacement at the same time point as extrapolated from creep curve before illumination. The average light-induced length reduction against four different drag forces is plotted in Figure 3c. Against a drag force of 1 pN, the rotary motor reduced the displacement length of the tethered particles by almost 12%. This corresponds to a work per bead around 100-200 pN·nm, which is in the expected range for a few motors^6^. Length reduction decreased with increasing flow forces. These force values appear rather low compared to the tens of pN expected to be involved in mechanotransduction by integrins^20^. However, entropic forces arising from stretching of coiled chains scale inversely with their persistence length and thus 50 times higher forces are expected to be applied by the twisted PEG5000 chains in the cell experiments, matching the relevant range for mechanical stimulation of cellular signals.

**Figure 3.**
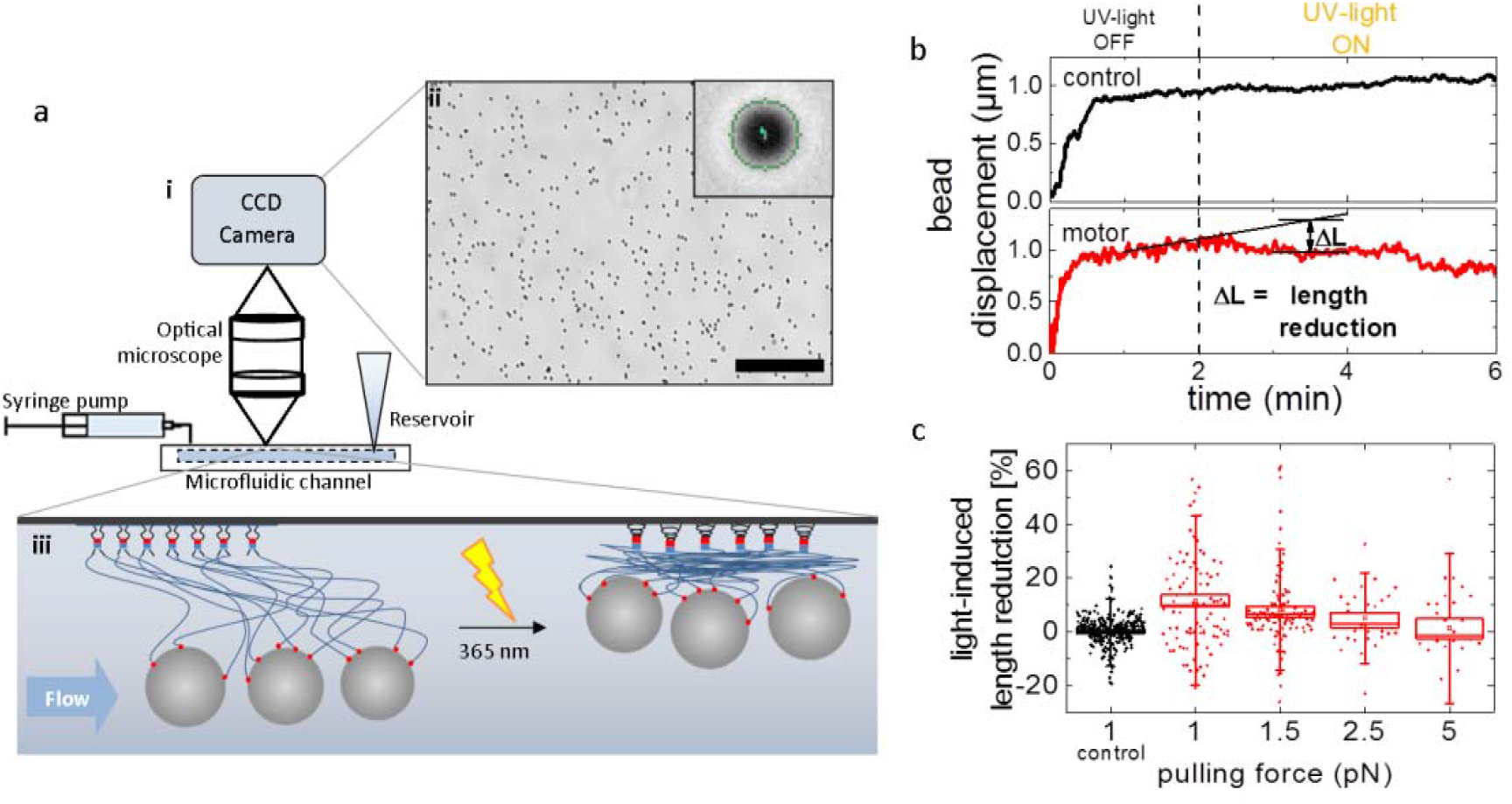
Measurement of forces applied by the light-driven rotary motor. (a,i) Experimental setup for the observation of tethered particle motion in a microfluidic flow channel. (a,ii) Optical microscopy determines the movement of hundreds of surface-tethered beads in parallel (scalebar: 100 μm); the inset demonstrates tracking of one exemplary 500 nm bead. (a,iii) Sketch of the molecular arrangement under flow before and after UV–light irradiation. Micro-beads are attached to the motor molecules via multiple DNA chains. Upon light activation, the molecular motor twists the DNA chains and the construct pulls the beads in. (b) Typical bead displacement versus time after start of flow with a drag force of 1 pN. The bead displacement is suddenly reduced after the molecular motor is activated by UV light. No such reduction is observed in control experiments with the static motor molecule. (c) Average light-induced length reduction after 2 min of UV-light irradiation for rotary motor molecule against different flow-induced drag forces and for the control with static molecules for 1 pN, normalized to the expected length of the construct (total n=564 beads).

Much of our understanding of the molecular mechanisms underlying force-stimulated cellular processes has been garnered from studies using micropipettes^21^, atomic force microscopes (AFM)^22^, or optical tweezers^23^ to apply forces to cells at subcellular level. The quantification of the applied forces, in parallel with the imaging of cell responses, have identified major players and force ranges in the mechanotransduction chain^24^. These physical methods, however, are limited in their ability to apply forces in multicellular or 3D environments, closer to physiological systems. Molecular approaches, like chemical-induced motors used by nature for intracellular force generation, enable a new dimension in mechanotransduction research. Salaita and coworkers ingeniously exploited a temperature-driven phase transition of a thin hydrogel film to pull on adhesive receptors of gel-bound cells^11^. This pioneering approach has, however, limitations for quantitative studies, as it does not allow gradual force regulation, and it requires encapsulation of the cell in a temperature-sensitive material, restricting its application in physiologically relevant contexts. Light-driven molecular machines inserted at engineered biointerfaces, as demonstrated in the current work, allow force application directly at molecular scale, without perturbation of other interface parameters or of an external probe or a pipette. The molecular character allows positional, magnitude, and time scale regulation of the applied force by tuning light localization and intensity. Its modular design allows flexible application to any receptor/ligand complex of choice without additional synthetic effort.

While this manuscript presents convincing demonstration of the potential of this tool, the development of the full potential of this method will motivate future work at different levels. More efficient motors to be activated at longer wavelengths^25^, understanding and quantification of rotation frequencies and force application rates and their modulation by the design of the flexible space are relevant open questions for improved performance. In addition, the study of force decay following illumination termination and its control through the relaxation mechanisms of the polymeric spacer offers additional ways to probe and understand the mechanical language in biological systems. Coupling to molecular force sensors (FRET tension probes^26 27 28^) could allow straightforward integration of force application and readout in simple additive steps. The flexibility of the design, not coupled to materials or special techniques, will allow extension of this technique to natural biointerfaces and *in vivo* contexts.

## METHODS

### 1. Preparation of ligand (or DNA)/motor/PEG/surface conjugates

#### 1.1 Preparation of motor/PEG/surface conjugate

N-hydroxysuccinimide (NHS) functionalized NEXTERION^®^ slide H (Schott, 1070936) was used as substrate for cell experiments. NEXTERION^®^ Slide H is coated with a cross-linked, multi-component non-fouling polymer layer and contains N-Hydroxysuccinimide (NHS) ester groups for direct covalent reaction with amine functionalized molecules.

The NEXTERION^®^ slide H was incubated with 50 μl of 1mM dibenzocyclooctyne-amine (DBCO-amine) solution in anhydrous DMSO for 1 h, and subsequently rinsed with Milli-Q water (200 μl) for 3 times. The substrate was then incubated with 50 μl of a 0.5 mM solution in Milli-Q water of polymer-motor conjugate (or control polymer-motor conjugate). After 12 h, the solution was removed and the substrate was then immersed in O-(2-Azidoethyl)-O’-methyl-triethylene glycol (PEG-azide) (20 mM in water, 50 μl) for 0.5 h to block the unreacted DBCO groups, and subsequently washed with Milli-Q water (200 μl) for 3 times. This substrate was immediately coupled to the biological ligand (next sections).

#### 1.2 Preparation of RGD/motor/PEG/surface conjugate

The motor/PEG/surface conjugate prepared as described in 1.1 was further incubated with cyclo(RGDfK-N_3_) solution (0.1 mg/mL in PBS, 50 μl, 0.1 mM) containing sodium ascorbate (1 mg/mL, 1 μl, 0.1 mM) and CuSO_4_·5H_2_O (1 mg/mL, 1 μl, 0.08 mM). The reaction was allowed to proceed for 6 h at room temperature. The modified substrate was rinsed with PBS (200 μl) for 3 times. The slides were used for cell experiments immediately after modification.

#### 1.3 Preparation of α-CD3/motor/PEG/surface conjugate

The motor/PEG/surface conjugate prepared as described in 1.1 was further incubated with biotin-PEG_3_-N_3_ solution (1 mg/mL in water, 50 μl, 2.2 mM) containing sodium ascorbate (1 mg/mL, 1 μl, 0.1 mM) and CuSO_4_·5H_2_O (1 mg/mL, 1 μl, 0.08 mM). The reaction was allowed to proceed for 3 h at room temperature and rinsed with Milli-Q water for 3 times. The substrate was then incubated with streptavidin (100 μg/mL in PBS, 50 μl) for 1 h at room temperature. After rinsed with Milli-Q water, the substrate was incubated with biotin antihuman CD3 Antibody (50 μg/mL in PBS, 50 μl) for 12 h at 4°C. The substrates were then rinsed with Ringer solution containing 0 mM Ca^2+^ (200 μl) for 3 times. The slides were used for cell experiments immediately after preparation.

#### 1.5 Preparation of α-CD28/motor/PEG/surface conjugate

The motor/PEG/surface conjugate prepared as described in 1.1 was further incubated with biotin-PEG_3_-N_3_ solution (1 mg/mL in water, 50 μl, 2.2 mM) containing sodium ascorbate (1 mg/mL, 1 μl, 0.1 mM) and CuSO_4_·5H_2_O (1 mg/mL, 1 μl, 0.08 mM). The reaction was allowed to proceed for 3 h at room temperature and rinsed with Milli-Q water for 3 times. The substrate was then incubated with streptavidin (100 μg/mL in PBS, 50 μl) for 1 h at room temperature. After rinsed with Milli-Q water, the substrate was incubated with biotin antihuman CD28 Antibody (50 μg/mL in PBS, 50 μl) for 12 h at 4°C. The substrates were then rinsed with Ringer solution containing 0 mM Ca^2+^ (200 μl) for 3 times. The slide was used for cell experiments immediately after preparation.

#### 1.6 Preparation of DNA/motor/PEG/surface conjugate

The microfluidic channel was constructed by sandwiching a double-sided polyiminde film (Kapton tape) between a previously modified motor/PEG NEXTERION^®^ slide H (as indicated in section 1.1) and a microscopy slide. A 1 mm x 10 mm rectangular channel was cut into the Kapton tape using a CO_2_ laser cutter. Two 0.8 mm holes were drilled into the microscopy slide as in- and outlet for solutions. The tubing was connected to the in- and outlet via 10 μl Pipette tips gently pushed into the holes.

The DNA-tethers were attached to the motor molecules in a two-step process. First, a DNA-oligomer was covalently attached to the motor molecules providing anchoring points for the longer DNA construct that serves as a tether for the micro-beads. The 50 bp, 3’ azide terminated single strand DNA-oligomer 5’-CATCACCTTGCTGAACCTCAAATATCAAACCCTCA ATCAATATCTGGTCA-3’) (Integrated DNA Technology) was covalently attached to the motor/PEG NEXTERION^®^ slide H by incubating the channel with a 10 μM DNA-oligomer solution in nuclease-free water for 12 h in presence of CuSO_4_·5H_2_O and Na-ascorbate. Then, the channel was washed three times with 10 μl nuclease-free water and the channel walls were passivated with 10 mg/ml Western Blocking Reagent (Sigma Aldrich) in PBS for 1 h. The blocking solution was replaced every 15 min. In the second step, a 2.7 kbp long DNA construct was attached to the surface wall via hybridization with the DNA-oligomers. The DNA construct was assembled from circular M13mp18 ssDNA (New England Biolabs) and functionalized with biotin to enable attachment of streptavidin-functionalized beads as described in Ref^22^. The region complementary to the DNA-oligomer was left single-stranded to enable efficient hybridization. The DNA solution (7 μl, 100 pM) was pipetted into the channel and incubated for 12 h. Subsequently, streptavidin-coated beads (Dynabeads MyOne C1, Thermo Fisher Scientific) were injected into the fluid channel for 5 min to attach to the DNA via specific binding of biotin and streptavidin. Previously, the beads were washed extensively and diluted to a concentration of 1 g/mL in PBS. After tethering, the chamber was flipped upside down and loose beads were washed out by applying a gentle fluid flow of 0.5 μl/min. The chamber was used immediately for experiments. The samples for the control experiments were prepared in the same way using the non-rotary motor (control).

### 2. Cell experiments

#### 2.1 Cell culture and reagents

Fibroblast L929 cell line were cultivated at 37 °C in 5 % CO_2_ in RPMI medium (Gibco) supplemented with 10% fetal bovine serum (Invitrogen) and 1 % P/S (Invitrogen). Cells were used between passages P4 and P16.

Mouse embryonic fibroblasts (MEFs) expressing paxillin-RFP were cultured in high-glucose DMEM (ThermoFisher) supplemented with 10% FBS (manufacturer), 2 mM GlutaMAX (Gibco) and 1% penicillin-streptomycin (manufacturer). For imaging experiments, the culture medium was replaced with phenol red-free DMEM Fluorobrite (Gibco), supplemented with the same supplements as above. For experiments with ROCK inhibitor Y27632 (10 μM), cells were pre-incubated for 30-60 minutes before imaging.

Jurkat T cells were cultured in RPMI-1640 medium (ThermoFisher) with 10% FBS (ThermoFisher) and 1% penicillin-streptomycin (Gibco). Primary human CD4^+^ T cells were isolated by untouched CD4^+^ T cell isolation kit (Miltenyi Biotec). Human CD4^+^ T cells were activated by Dynabeads Human T-Activator CD3/CD28 (ThermoFisher) and cultured in AIMV medium (ThermoFisher) with 10% FBS (ThermoFisher) and in presence of 33 U/ml human IL-2 (premium-grade, Miltenyi Biotec).

#### 2.2 Optimization of illumination conditions in MEFs

A UGA-42 Firefly point scanning laser device (RAPP Optoelectronic) with a 365 nm laser was coupled to an inverted epifluorescence Zeiss Axio Observer Z1 microscope. The laser focus was calibrated using a chromium calibration slide (RAPP Optoelectronic) prior to each experiment. At 50% duty cycle (the power used for the cell experiments and the minimal output we could measure using the laser meter), the power of the laser was 660 nW (measured using a laser meter from LABMASTER^®^) with a laser spot size of 4.4 μm^2^ with the usage of an OD3 filter. This results in an irradiance of 15 mW/cm^2^ (OD2: 150 mW/cm^2^). The substrate underneath the cell was illuminated using a custom circular ROI with a diameter of 8.86 μm (Area= 62 μm^2^) by using the SysCon software. The ROI was scanned with 5 scans per illumination (Duration 234.75 ms; 5 runs/object 1234.75 ms), repeated every 20s for duration of 15 min. The sample was irradiated simultaneously while acquiring RFP images for widefield fluorescence microscopy.

Initial exposure experiments were performed with a neutral density (ND) filter of OD2 (1% transmission) resulting in a light dose of 150 mW/cm^2^, with a cyclical illumination regime of one illumination pulse (40 scans per illumination) per 20s (Supplemental Fig. 3). All cells monitored showed immediate cell retraction (within 6 minutes) of the illuminated cell areas, on both motor substrate (8 out of 8) and control substrate (4 out of 4). Cells did not show this behavior when lowering the laser transmission by using a stronger ND filter (OD3 – 0.1% transmission), although cell retraction was still observed on motor substrates (86%, 6 out of 7 cells) and control substrates (60%, 3 out of 5). Only when additionally lowering the amount of scans per illumination pulse (5 scans per illumination) did most cells on both motor (75%, 9 out of 12) and control substrates (80%, 8 out of 10) not show any cell retraction. These illumination conditions were used for the experiments.

#### 2.3 Live-cell imaging for focal adhesions

Cells expressing intermediate to low levels of paxillin were chosen to avoid overexpression artifacts. Cells were imaged on an inverted epifluorescence Zeiss Axio Observer Z1 microscope using a Plan-Apo 40x Oil Objective (NA=1.4) and 1.6 Tubelens, coupled to an Axiocam 506 CCD camera. Samples were illuminated using the Colibri 530 nm LED coupled with the DsRed filterset (set 43) with 8% power of 530 nm LED for 500 ms exposure time for each time point. In addition, brightfield images were taken (100 ms exposure) with the TL lamp set to 3.5V. Images were obtained with 2×2 binning and 2x analog gain every 30s for 15 minutes. Images were captured using the Zen

#### 2.4 Focal adhesion images processing and analysis

All image processing was done using the Fiji distribution of ImageJ^29^ with the Morpholib plugin library^30^ For the reference images (Fig 2a and b) and Supplemental Movie 1, paxillin-RFP images were first filtered with a Gaussian Blur (sigma=1), then the Grays LUT was inverted and contrast was enhanced.

To analyze illuminated versus non-illuminated areas, first ROIs containing several FAs for these areas were cropped, and then each ROI was separately processed and analyzed. For Focal Adhesion segmentation a custom ImageJ macro was developed. In short, images were first filtered using a Gaussian blur (sigma=1.5), then a 3D White top Hat filter (element=cube, xyz-radii were all set at 5) from the Morpholib plugin was applied, after which background was subtracted using a rolling ball of 20. Then a mask was generated using the Otsu autothreshold. Small noise particles were then excluded by Area opening (pixel size=20). The total size of all FAs was then measured using the resulting mask. The relative increase in total FA area was determined as total FA size at t/ total FA size at t_0_.

#### 2.5 Optimization of irradiation conditions in T cells

For T cell activation, we employed a Zeiss Cell Observer HS system with a 20 × alpha objective and an AxioCam MRm Rev. 3. The DAPI channel (100% power, 10 mW/cm^2^) was used for UV irradiation. A sequence of ten UV pulses with a duration of 1, 0.5 or 2 seconds were applied within 1 minute to activate rotation of the motor. The Ca^2+^ fluorescent signal was followed for 15 – 20 min.

#### 2.6 Calcium imaging and data analysis

Jurkat T cells and primary T cells were loaded with Fluo-4/AM (1 μM) in serum free RPMI-1640 media at room temperature for 30 min. Afterwards cells were pelleted by centrifugation, followed by one wash using Ringer’s solution containing 0 mM Ca^2+^. Then cells were settled on the motor-linked hydrogels for 8 min before measurement. Afterwards, the cells were washed once with Ringer’s solution containing 0 mM Ca^2+^. Then the same volume of Ringer’s solution containing 2 mM Ca^2+^ was added prior to the start of calcium imaging. 15 min later, UV illumination was conducted for 1 min followed with another 15 min of imaging. To detect a general capability of the cells to induce Ca^2+^ influx, 1 μM thapsigargin is added 10 min after UV illumination.

For calcium imaging, we employed a Zeiss Cell Observer HS system with a 20 × alpha objective and an AxioCam MRm Rev. 3. During the experiment, the fluorescence of Fluo-4 was acquired with a 38HE filter set every 5 sec. Mean fluorescence intensity of Fluo-4 was quantified with ImageJ. The cell spreading areas were determined using the Wand (tracing) Tool in ImageJ and then quantified with ImageJ. The peak of Ca^2+^ influx (ΔPeak) is the maximum relative the fluorescence of Fluo-4 (normalized to base line before UV illumination) from first UV pulse to 120 s.

### 3. Tethered particle motion experiments

#### 3.1 Particle tracking in the flow cell

All Flow Cell measurements were performed with a syringe pump (AL-1000 World Precision Instruments, Saratosa, USA) equipped with a 3 ml syringe (BD Diagnostics). A CCD camera (ImagingSource, Bremen, Deutschland) with 3.072×2.048 pixels mounted on a standard optical microscope with a 50x magnification was used for recording the movement of surface-tethered beads in one field of view. Prior to every experiment, the zero position of the beads was determined by recording the Brownian motion of the beads for 60 s with a sampling rate of 5 fps. Subsequently, a constant flow of 2 μl/min to 15 μl/min was applied for 8 minutes. After 2 min of constant flow, the UV LED with a maximum intensity at a wavelength of 365 nm and a power of ~0.75 W/cm^2^ was switched on for 4 minutes. Force calibration was performed in DNA overstretching experiments with linear flow ramps from 0 μl/min to 1000 μl/min controlled via the syringe pump. The actual flow velocity was monitored via the height of the fluid in the reservoir (Figure 3a).

#### 3.2 Data analysis

The digital videos were analyzed using the open source software *ImageJ 1.50i* (Wayne Rasband, National Institute of Health, USA) and a *Particle Tracker* plugin for *ImageJ* written by Sbalzarini and Koumoutsakos. The *Particle Tracker* provides *x* and y positions of each individual bead for every frame. The *ImageJ* software includes all trajectories into a table that was further analyzed in MATLAB regarding their Brownian motion and bead displacement under fluid flow. Before data analysis, all trajectories were drift-corrected using a master curve generated by averaging the trajectories of three to five immobile beads^31^. The heterogeneity of the molecular motor attachment results in a statistical distribution of biotin binding sites leading to variety of tethered particle motion. The Brownian motion of all beads in one frame was recorded for 60s prior to every measurement. Only trajectories with a RMS position fluctuation above 100 nm were included in further analysis to omit unspecific beads with non-specific attachment. This lower limit reflects an effective tether length L=(3*(RMS)^2^)/2b^32^ of above 375 nm assuming a Kuhn-length of *b* =40 nm.

## Data Availability Statement

The authors declare that all data supporting the findings of this study are available within the paper and the supplementary information.

## Acknowledgements

A.d.C., B.Q. and M. H. acknowledge support from Deutsche Forschungsgemeinschaft (SFB 1027). M. H. acknowledges support from the Alexander von Humboldt Foundation and Deutsche Forschungsgemeinschaft (SPP 1782). A.d.C. acknowledges support from European Union (FET Mechanocontrol). N.G. acknowledges support from the Laboratory of Excellence for Complex Chemical Systems (LabEx CSC). A.G. acknowledges support from the US National Institute of Health (R01 EB024322). We thank Dr. Emmanuel Terriac and Dr. Aleeza Farrukh for their assistance with imaging.

## Author Contributions Author Contributions

The idea to use the molecular motor to apply force to individual cellular receptors was suggested by A.d.C., and refined in discussions with N.G., R. B. and A.G.. The polymer-motor conjugates were designed by N. G. and synthesized by J.R.C.I. and D.D.. A.d.C. and Y.Z. designed the biointerfaces. Y.Z. developed the motor-substrates for all the experiments. J.Z. performed the first studies with T cells together with Y.Z.. A.d.C., A.G. and M.H. conceived the experiments with the fibroblasts. M.H. carried out the experiments with fibroblasts and quantified the results. D.Z. engineered the RFP-paxillin fibroblasts. The experiments with T cells were designed by B.Q., M.H. and R.Z. and were carried out by R.Z.. J. B. and R.B. implemented the tethered particle method to quantify motor applied forces. All authors contributed to manuscript writing and/or correction.

## Supplementary Information

is available for this paper.

## Correspondence and requests for materials

should be addressed to aranzazu.delcampo@leibniz-inm. de.

## References

1 Humphrey, J. D., Dufresne, E. R. & Schwartz, M. A. Mechanotransduction and extracellular matrix homeostasis. Nature Reviews Molecular Cell Biology 15, 802, doi:10.1038/nrm3896 (2014).

2 Matthews, B. D., Overby, D. R., Mannix, R. & Ingber, D. E. Cellular adaptation to mechanical stress: role of integrins, Rho, cytoskeletal tension and mechanosensitive ion channels. Journal of Cell Science 119, 508, doi:10.1242/jcs.02760 (2006).

3 Schliwa, M. & Woehlke, G. Molecular motors. Nature 422, 759–765, doi:10.1038/nature01601 (2003).

4 Dattler, D. et al. Design of Collective Motions from Synthetic Molecular Switches, Rotors, and Motors. Chem Rev 120, 310–433, doi:10.1021/acs.chemrev.9b00288 (2020).

5 Koumura, N., Zijlstra, R. W. J., van Delden, R. A., Harada, N. & Feringa, B. L. Light-driven monodirectional molecular rotor. Nature 401, 152–155, doi:10.1038/43646 (1999).

6 Li, Q. et al. Macroscopic contraction of a gel induced by the integrated motion of light-driven molecular motors. Nat Nanotechnol 10, 161–165, doi:10.1038/nnano.2014.315 (2015).

7 Foy, J. T. et al. Dual-light control of nanomachines that integrate motor and modulator subunits. Nature Nanotechnology 12, 540–545, doi:10.1038/nnano.2017.28 (2017).

8 Colard-Itté, J.-R. et al. Mechanical behaviour of contractile gels based on light-driven molecular motors. Nanoscale 11, 5197–5202, doi:10.1039/C9NR00950G (2019).

9 Baroncini, M., Silvi, S. & Credi, A. Photo- and Redox-Driven Artificial Molecular Motors. Chem Rev 120, 200–268, doi:10.1021/acs.chemrev.9b00291 (2020).

10 Sniadecki, N. J. et al. Magnetic microposts as an approach to apply forces to living cells. Proceedings of the National Academy of Sciences 104, 14553–14558 (2007).

11 Liu, Z. et al. Nanoscale optomechanical actuators for controlling mechanotransduction in living cells. Nat Methods 13, 143–146, doi:10.1038/nmeth.3689 (2016).

12 Riveline, D. et al. Focal Contacts as Mechanosensors. The Journal of Cell Biology 153, 1175 (2001).

13 Liu, B., Chen, W., Evavold, B. D. & Zhu, C. Accumulation of dynamic catch bonds between TCR and agonist peptide-MHC triggers T cell signaling. Cell 157, 357–368, doi:10.1016/j.cell.2014.02.053 (2014).

14 Kim, S. T. et al. The αβ T Cell Receptor Is an Anisotropic Mechanosensor. Journal of Biological Chemistry 284, 31028–31037, doi:10.1074/jbc.M109.052712 (2009).

15 Liu, Y. et al. DNA-based nanoparticle tension sensors reveal that T-cell receptors transmit defined pN forces to their antigens for enhanced fidelity. Proceedings of the National Academy of Sciences 113, 5610–5615 (2016).

16 Judokusumo, E., Tabdanov, E., Kumari, S., Dustin, M. L. & Kam, L. C. Mechanosensing in T lymphocyte activation. Biophys J 102, L5–7, doi:10.1016/j.bpj.2011.12.011 (2012).

17 Bashour, K. T. et al. CD28 and CD3 have complementary roles in T-cell traction forces. Proceedings of the National Academy of Sciences 111, 2241, doi:10.1073/pnas.1315606111 (2014).

18 Garcia-Lopez, V. et al. Molecular machines open cell membranes. Nature 548, 567–572, doi:10.1038/nature23657 (2017).

19 Yang, D., Ward, A., Halvorsen, K. & Wong, W. P. Multiplexed single-molecule force spectroscopy using a centrifuge. Nature Communications 7, 11026, doi:10.1038/ncomms11026 (2016).

20 Sun, Z., Costell, M. & Fässler, R. Integrin activation by talin, kindlin and mechanical forces. Nature Cell Biology 21, 25–31, doi:10.1038/s41556-018-0234-9 (2019).

21 Oesterhelt, F., Rief, M. & Gaub, H. E. Single molecule force spectroscopy by AFM indicates helical structure of poly(ethylene-glycol) in water. New Journal of Physics 1, 6–6, doi:10.1088/1367-2630/1/1/006 (1999).

22 Koussa, M. A., Halvorsen, K., Ward, A. & Wong, W. P. DNA nanoswitches: a quantitative platform for gel-based biomolecular interaction analysis. Nat Methods 12, 123–126, doi:10.1038/nmeth.3209 (2015).

23 Kadem, L. F. et al. High-Frequency Mechanostimulation of Cell Adhesion. Angewandte Chemie International Edition 56, 225–229, doi:10.1002/anie.201609483 (2017).

24 Jaalouk, D. E. & Lammerding, J. Mechanotransduction gone awry. Nature Reviews Molecular Cell Biology 10, 63, doi:10.1038/nrm2597 (2009).

25 Roke, D., Wezenberg, S. J. & Feringa, B. L. Molecular rotary motors: Unidirectional motion around double bonds. Proceedings of the National Academy of Sciences 115, 9423, doi:10.1073/pnas.1712784115 (2018).

26 Polacheck, W. J. & Chen, C. S. Measuring cell-generated forces: a guide to the available tools. Nature Methods 13, 415, doi:10.1038/nmeth.3834 (2016).

27 Grashoff, C. et al. Measuring mechanical tension across vinculin reveals regulation of focal adhesion dynamics. Nature 466, 263–266, doi:10.1038/nature09198 (2010).

28 Stabley, D. R., Jurchenko, C., Marshall, S. S. & Salaita, K. S. Visualizing mechanical tension across membrane receptors with a fluorescent sensor. Nat. Methods 9, 64–67, doi:10.1038/nmeth.1747 (2012).

29 Schindelin, J. et al. Fiji: an open-source platform for biological-image analysis. Nat Methods 9, 676–682, doi:10.1038/nmeth.2019 (2012).

30 Legland, D., Arganda-Carreras, I. & Andrey, P. MorphoLibJ: integrated library and plugins for mathematical morphology with ImageJ. Bioinformatics 32, 3532–3534, doi:10.1093/bioinformatics/btw413 (2016).

31 Nelson, P. C. et al. Tethered Particle Motion as a Diagnostic of DNA Tether Length. The Journal of Physical Chemistry B 110, 17260–17267, doi:10.1021/jp0630673 (2006).

32 Sindilariu, P.-D., Brinker, A. & Reiter, R. Waste and particle management in a commercial, partially recirculating trout farm. Aquacultural Engineering 41, 127–135, doi:10.1016/j.aquaeng.2009.03.001 (2009).

