## Extended Data for "Optoregulated force application to cellular receptors using molecular motors"

#### Preparation of ligand (or DNA)/motor/PEG/surface conjugates

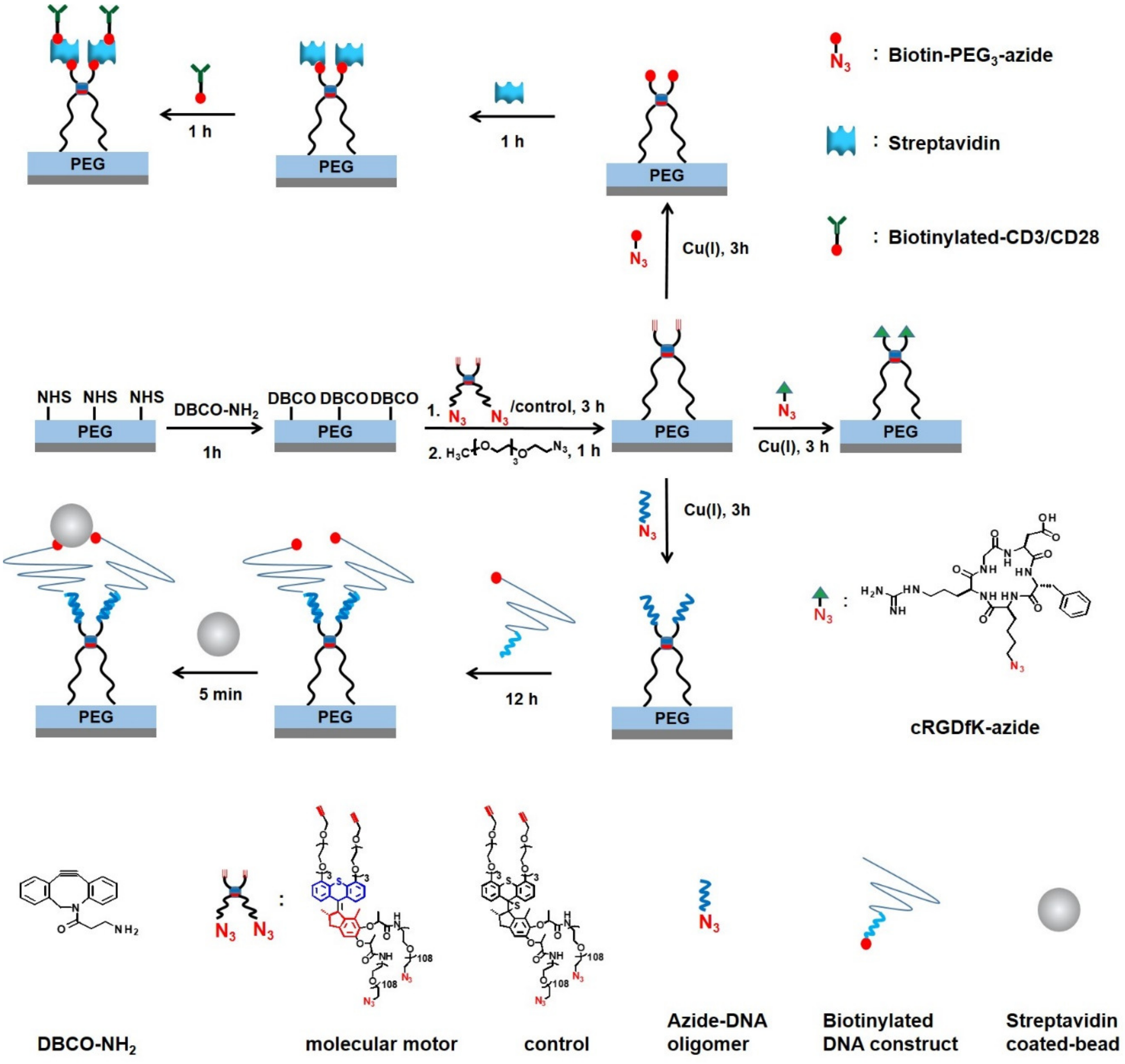


Supplementary Scheme 1. Schematic illustration of the preparation of ligand (or DNA)/motor/PEG/surface conjugates through a multiple-step procedure.

#### Demonstration of specific functionalization of RGD/motor/PEG/surfaces

In order to demonstrate the specific and functional coupling of the RGD ligand to the motor/PEG/surface, cell adhesion experiments were performed. L929 fibroblasts were seeded on the motor/PEG/surface and on the RGD/motor/PEG/surface conjugates (Supplementary Figure 1). No cell attachment was observed on the motor/PEG/surface (Supplementary Figure 1a) or on the substrates incubated with RGD in absence of the motor (Supplementary Figure 1b). Cells attached and spread on RGD/motor/PEG/surfaces and showed normal morphology on the substrates over 7 days (Supplementary Fig. 1c), indicating that the spacer-motor conjugate was not toxic to the cells The density of attached and spread cells increased with the RGD density on the surface, which was regulated by the incubation concentration of RGD ligand in the surface modification step (Supplementary Fig. 2). Cells did not attach to substrates modified with the negative control peptide, RDG/motor/PEG/surface (Supplementary Fig. 1d). These results demonstrate that (i) the coupling reaction between motors and RGD is specific, (ii) only RGD mediates cell binding to the substrates, and (iii) binding of RGD ligand to the surface is mediated solely by the motor conjugate.

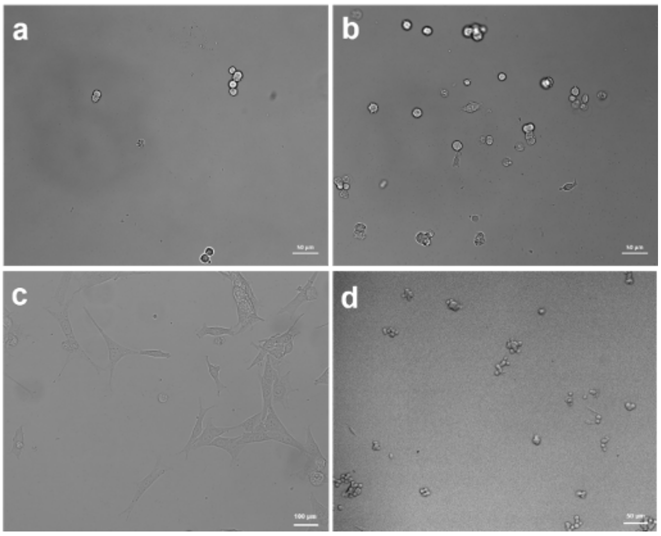


**Supplementary Figure 1:** L929 fibroblasts incubated on (a) motor/PEG/surface and (b) on substrates incubated with RGD without the motor for 24 hours. (c) Fibroblasts incubated on RGD/motor/PEG/surface after 7 days. (d) Fibroblasts incubated on RDG/motor/PEG/surface for 24 hours.

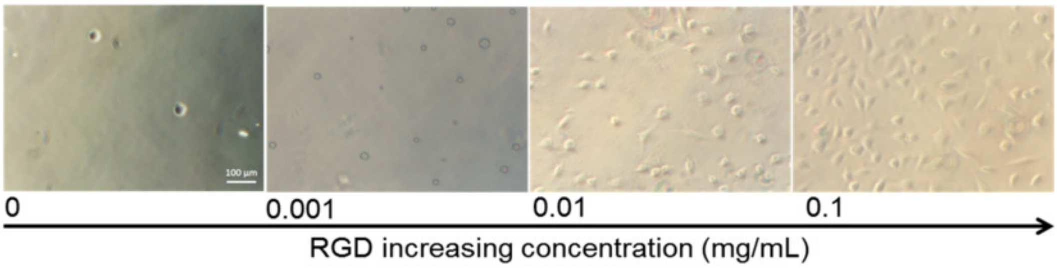


**Supplementary Figure 2:** Representative bright field images of L929 cells cultured on RGD/motor/PEG/surfaces at increasing concentrations of RGD peptide with a seeding number of 4000 cells/cm$^2$ after incubation for 24 hours.

#### Focal adhesion analysis

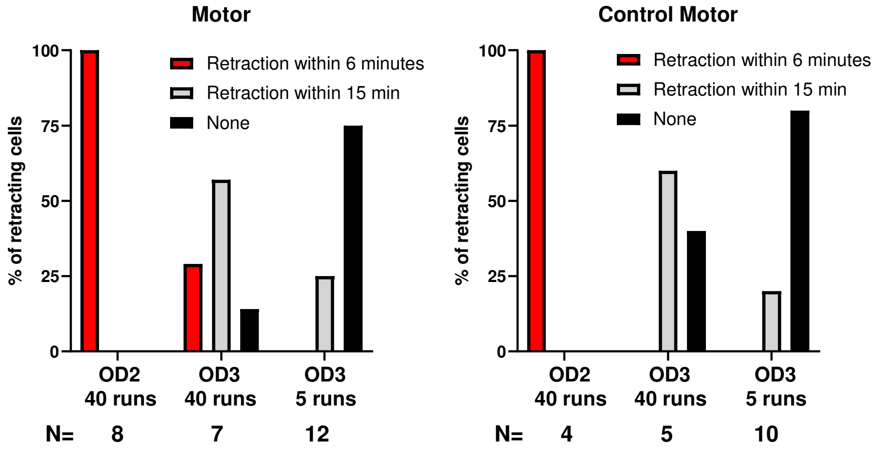


**Supplemental Figure 3.** Increasing UV-illumination dosage leads to cell retraction. Motor substrates (left) or control motor substrates (right) were illuminated with a scanning UV-laser (365 nm – 5 runs per 20s). Cell areas within the field of illumination were analyzed for cell retraction (removal of cell arm from illuminated area). OD is a neutral density filter blocking light transmission form the laser source, with OD2 and OD3 resulting in 1% and 0.1% light transmission respectively. N represents the number of cells analyzed for each condition.

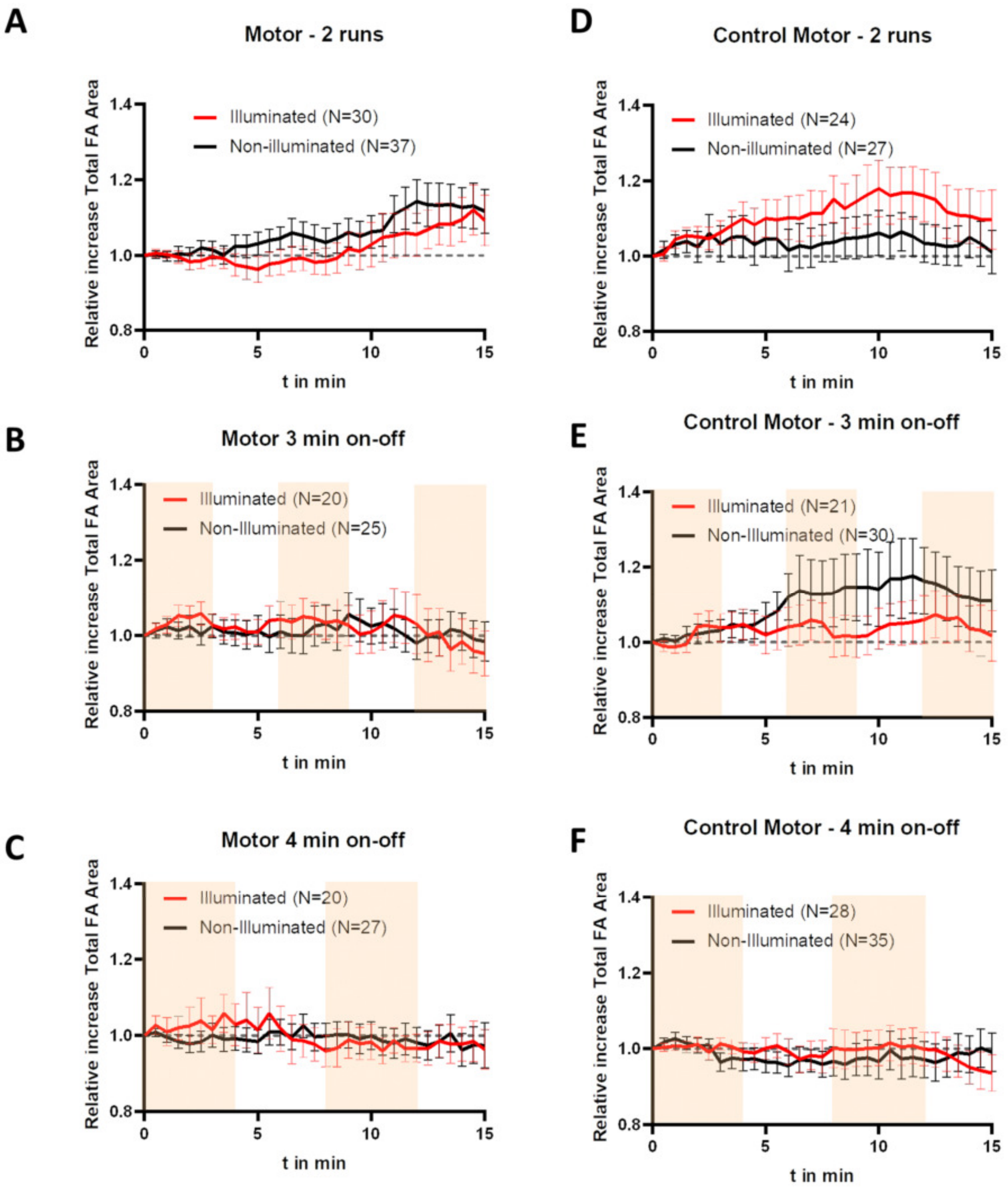


**Supplemental Figure 4.** Control experiments. Plots represent the mean ± s.e.m. of relative increase in Total Focal Adhesion Area of Focal adhesions analyzed within N# of ROIs of cells seeded on motor (a-c) or control motor (d-f) substrates irradiated with different irradiation programs. (a,d) 2 runs per 20s irradiation. (b,e) 5 runs per 20s -3 min on – 3 min off. Illuminated times are marked in orange. (c,f) 5 runs per 20s - 4 min on – 4 min off. Illuminated times are marked in orange. All data was generated from three independent experiments, except (c) which was generated from two independent experiments.

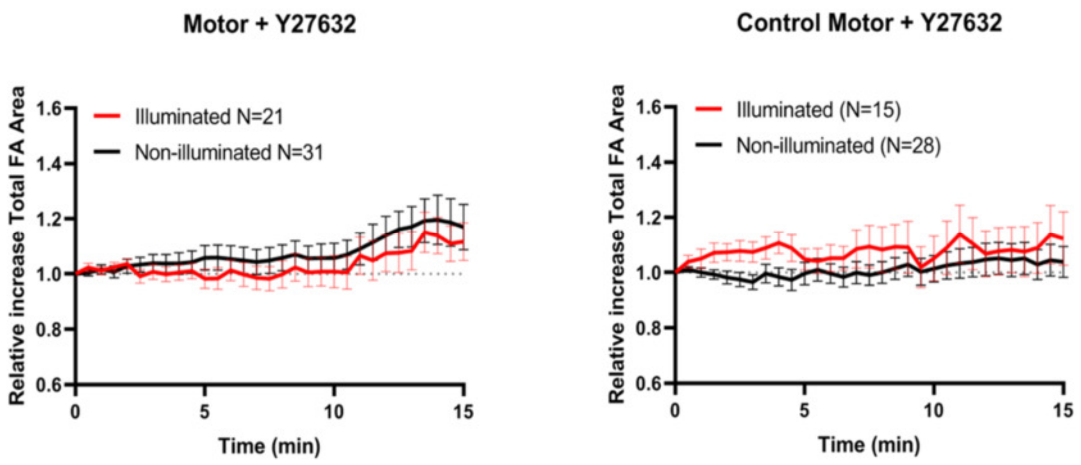


**Supplemental Figure 5.** ROCK inhibition perturbs motor-induced FA activation. Cells were pre-incubated with ROCK inhibitor Y27632 (10 µM) for 30-50 min. Plots represent the mean ± s.e.m. of relative increase in Total Focal Adhesion Area of Focal adhesions analyzed within N# of ROIs of 21 cells from three independent experiments seeded on motor (left) or of 15 cells from two independent experiments on control motor (right) substrates.

#### Literature analysis of FA responses to applied forces with physical devices.

Previous studies exerting extracellular force on a single molecule level, such as magnetic bead pulling experiments with RGD-coated beads, induced adhesion reinforcement within seconds, i.e. when 3s 130 pN force pulses were applied in an oscillatory fashion (6 pulses with 4s relaxation times)[1]. Similarly, the high-frequency oscillatory push-pulling-force of light-activated RGD-coupled azobenzene-PEG substrates increased the force needed to detach cells coupled to an AFM cantilever on seconds scale[2]. In contrast, force-induced FA growth has mostly been observed on the scale of minutes. Low nN-scale forces applied to integrins on the basal side of the cell using PDMS microposts containing magnetic nanowires, resulted in a 25% increase in relative FA size (as measured by vinculin-GFP recruitment) when a 10 min sustained magnetic field was applied. The observed increase was even higher (75%) when the force was applied in an oscillatory fashion (2 min on-off states for 10 minutes). It should be noted however that this method causes relatively large deformations on a cellular level. Interestingly, even when 13-50 pN forces were applied on a more molecular scale, using polymer particles coupled to RGD which shrink under influence of near-infrared light, FA growth occurred in the order of tens of minutes[3]. Application of a 10 Hz oscillatory force increased FA size (as monitored by paxillin recruitment) to 5 times the initial area. This mirrors the results of our method, which also acts on a single molecule scale and also induced FA growth in a similar time-scale.

#### Additional experiments with T cells

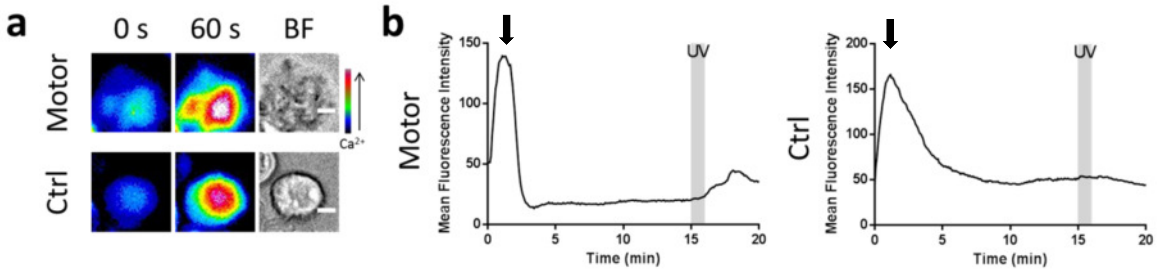


**Supplementary Figure 6.** Activation of Fluo-4-AM loaded Jurkat T cells on the αCD3/motor/PEG/surface. Control experiments using the non-rotary motor were also run in parallel. The cells were incubated in 0 mM $Ca^{2+}$ Ringer's solution at room temperature for 8 min. Then the same volume of 2 mM $Ca^{2+}$ Ringer's solution were added (0 min). At the time point of 15 min, the cells were illuminated by UV. A sequence of ten UV pulses with a duration of 1 second were applied within 1 minute and Fluo-4 fluorescent signal was followed for 20 mins. (a) Heat map of Fluo-4 fluorescence intensity. BF stands for bright field. Scale bars are 5 μm. (b) Analysis of intracellular $Ca^{2+}$ dynamics in Jurkat T cells shown in (a).

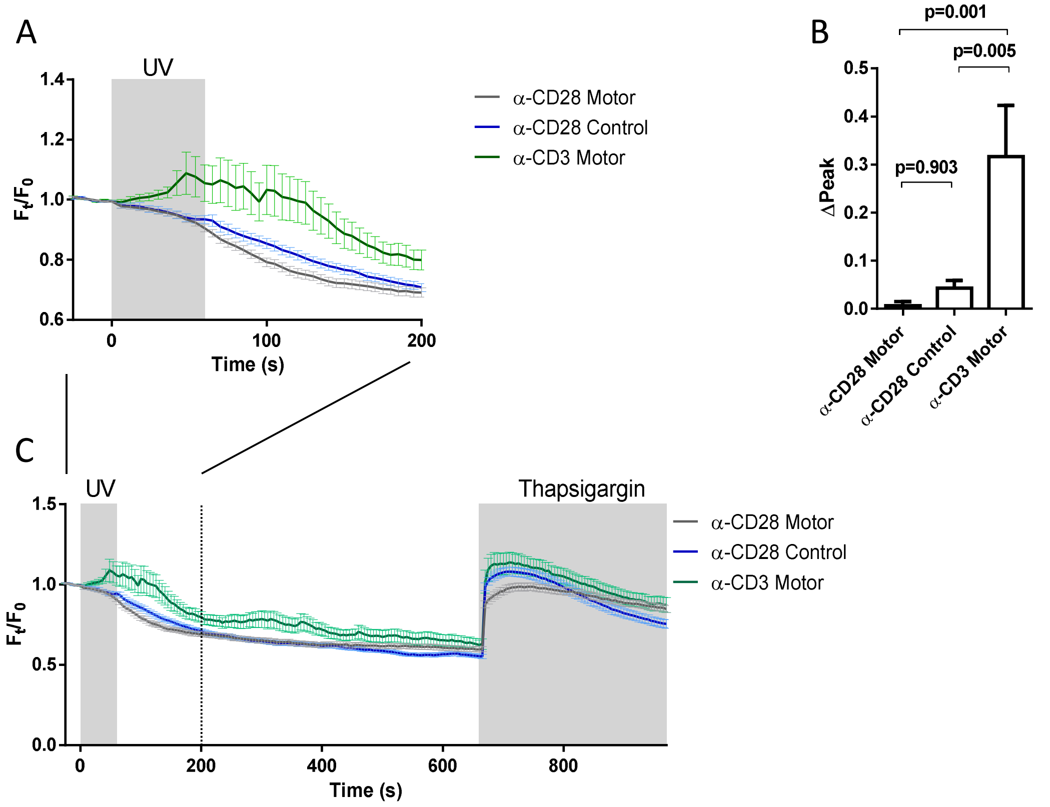


**Supplementary Figure 7.** Response of T cells on αCD28/motor/PEG/surfaces and controls. Jurkat T cells loaded with Fluo-4-AM were seeded on the substrate for 15 min before UV illumination. 10 pulses (1 s duration) of UV light were applied for a total duration of 1 min. (a) αCD28/motor does not induce $Ca^{2+}$ influx. αCD3/motor serves as a positive control. (b) Quantification of the maximum $Ca^{2+}$ influx (ΔPeak) in a. (c) UV illumination does not hamper the capacity of $Ca^{2+}$ influx. 10 min After UV illumination, the cells were activated by thapsigargin (1 µM) to induce the maximum $Ca^{2+}$ influx. The results were from 4 independent experiments (α-CD28 Motor, n=115 cells; α-CD28 Control, n=120 cells; α-CD3 Motor, n=120 cells) and represented as mean ± s.e.m.

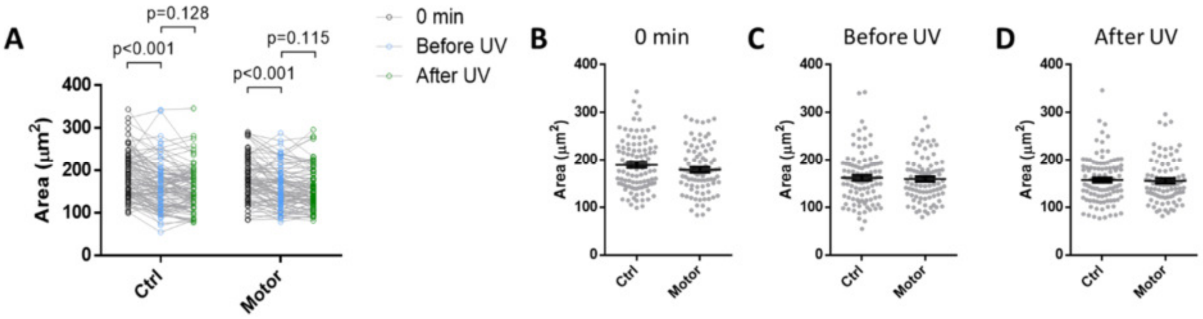


**Supplementary Figure 8.** Mechanical force applied by the motor does not change the contact area at the IS. Fluo-4 loaded Jurkat cells were seeded on αCD3 modified substrates. The CD3 antibody was linked with the surface via non-rotary motor (control, 100 cells) or rotary motor (88 cells). The contact areas were determined at three time points: the beginning of measurement (0 min), before and after UV illumination. (a) The contact area at the IS does not change upon rotation of the motor. Paired t-test was used for the statistics. (b-d) No difference in the contact area is found between control and motors. The results were from 6 independent experiments and represented as mean ± s.e.m..

#### Force measurements by observation of tethered particle motion

##### Creep in tethered particle motion

After onset and stabilization of flow (1 min), the tethered beads are typically displaced by a few hundred nanometers. The variation in initial displacement can be explained by a variation of the exact number of tethers and by the distribution of tether attachment on the Nexterion substrate. We then observed creep of the beads position, i.e. beads were continuously displaced in the flow direction. We attribute the creep to relaxation processes in the substrate gel and to a detachment and reattachment of the non-covalent biotin-streptavidin bonds of DNA tethers to the bead surfaces.

##### Method and scaling arguments

In the cell experiments, motor-chain conjugates were coupled to a Nexterion PEG-hydrogel, resulting in a random distribution of effective tether length between receptors and surface. The heterogeneous distribution of tether lengths is expected to generate a distribution of pulling forces applied to cellular focal adhesions. We introduce the video microscopy of tethered particle motion (TPM)[4] as a novel approach that allows investigation of hundreds of different motor-chain conjugates in parallel. Under flow and light exposure, this method allows parallel observation of drag force-induced displacement and light-induced retraction of hundreds of beads by optical microscopy. Other single molecule experiments such as atomic force microscopy (AFM) or optical or magnetic tweezers require the statistical evaluation of many repetitions to achieve piconewton force resolution. The time necessary to link single molecules and to record repeated force measurements impedes the investigation of heterogeneous systems, where many locations have to be probed to determine the distribution of mechanical properties.

In our TPM experiment, molecular motors are attached to the surface of the channel using the same PEG hydrogel functionalization as in the cell experiments. Beads with a diameter of 500 nm, comparable to the size of a focal adhesion, are tethered to the molecular motors by DNA chains with a length of 1.7 µm. The beads are connected to a small number of tethers, which resemble the attachment of focal adhesions by multiple motor-chain conjugates in the cell experiment. We use a DNA construct[5] which offers great flexibility in the attachment of functional end groups for linking the chains.

The observation of motion of tethered microparticles by video microscopy requires longer spacer chains than the ones used in the cell experiments. The contour length of the DNA construct (1700 nm) is about 53 times that of the $PEG_{5000}$ linkers (32 nm) in the cell experiment. The persistence length of the dsDNA (20 nm) is about 57 times that of $PEG_{5000}$ chains (0.35 nm[6]). Thus, the overall coiling geometry the two experiments is scaled and we can expect a similar relative reduction in extension upon twisting of polymer pairs in the entropic low-force regime. In absolute numbers, the same rotational twist induced by the motor molecule will induce a 50 times smaller length reduction in the 50 times shorter PEG chains, but this relative extension will produce a 50 times larger force than in the DNA experiments as all entropic forces for the same relative polymer extension scale as $k_B T/P$, where $P$ is the persistence length[7]. This scaling ultimately needs confirmation for the case of twisted pairs of polymer chains by an adequate model or simulations.

The dsDNA chains are constructed by hybridizing an ssDNA with a set of matching oligomers with a length of 60 bp each. The persistence length of this construct was determined by fitting an extensible worm-like chain model to the extension-force curves of single dsDNA constructs. The extension-force curves were recorded by analyzing the displacements of tethered beads in an increasing flow. The value of 20 nm for the persistence length is significantly lower than the values around 50 nm typically reported for dsDNA[8]. We attribute the difference to the high number of nicks every 60 base pairs, i.e. of breaks in the phosphate backbone of one strand of the dsDNA.

### References


1 Matthews, B. D., Overby, D. R., Mannix, R. & Ingber, D. E. Cellular adaptation to mechanical stress: role of integrins, Rho, cytoskeletal tension and mechanosensitive ion channels. *Journal of Cell Science* **119**, 508, doi:10.1242/jcs.02760 (2006).

2 Kadem, L. F. *et al.* High-Frequency Mechanostimulation of Cell Adhesion. *Angewandte Chemie International Edition* **56**, 225-229, doi:10.1002/anie.201609483 (2017).

3 Liu, Z. *et al.* Nanoscale optomechanical actuators for controlling mechanotransduction in living cells. *Nat Methods* **13**, 143-146, doi:10.1038/nmeth.3689 (2016).

4 Nelson, P. C. *et al.* Tethered Particle Motion as a Diagnostic of DNA Tether Length. *The Journal of Physical Chemistry B* **110**, 17260-17267, doi:10.1021/jp0630673 (2006).

5 Koussa, M. A., Halvorsen, K., Ward, A. & Wong, W. P. DNA nanoswitches: a quantitative platform for gel-based biomolecular interaction analysis. *Nat Methods* **12**, 123-126, doi:10.1038/nmeth.3209 (2015).

6 Oesterhelt, F., Rief, M. & Gaub, H. E. Single molecule force spectroscopy by AFM indicates helical structure of poly(ethylene-glycol) in water. *New Journal of Physics* **1**, 6-6, doi:10.1088/1367-2630/1/1/006 (1999).

7 Petrosyan, R. Improved approximations for some polymer extension models. *Rheologica Acta* **56**, 21-26, doi:10.1007/s00397-016-0977-9 (2017).

8 Bustamante, C., Smith, S. B., Liphardt, J. & Smith, D. Single-molecule studies of DNA mechanics. *Curr Opin Struct Biol* **10**, 279-285, doi:10.1016/s0959-440x(00)00085-3 (2000).
